# Comparative assessment of electronic nicotine delivery systems aerosol and cigarette smoke on endothelial cell migration: the Replica Project

**DOI:** 10.1101/2021.12.23.473979

**Authors:** Massimo Caruso, Rosalia Emma, Alfio Distefano, Sonja Rust, Konstantinos Poulas, Antonio Giordano, Vladislav Volarevic, Konstantinos Mesiakaris, Silvia Boffo, Aleksandar Arsenijevic, Georgios Karanasios, Roberta Pulvirenti, Aleksandar Ilic, Angelo Canciello, Pietro Zuccarello, Margherita Ferrante, Riccardo Polosa, Giovanni Li Volti, the Replica Project Group

## Abstract

Cigarette smoking is associated with impairment of repair mechanisms necessary for vascular endothelium homeostasis. Reducing the exposure to smoke toxicants may result in the mitigation of the harmful effect on the endothelium and cardiovascular disease development. Previous investigations performed by the tobacco industries evaluated in vitro the effect of electronic cigarette (e-cig) compared to cigarette smoke demonstrating a significant reduction in endothelial cell migration inhibition following e-cig aerosol exposure. In the present study, we replicated one of these studies, evaluating the effects of cigarette smoke on endothelial cell migration compared to e-cig and heated tobacco products. We used a multi-center approach (ring-study) to verify the robustness and reliability of the results obtained in the replicated study. Consistently with the original study, we observed a substantial reduction of the effects of e-cig and tobacco heated products on endothelial cell migration compared to cigarette smoke. In conclusion, our study further confirms the importance of e-cig and tobacco heated products as a possible harm reduction strategy for cardiovascular diseases development in smokers.

**Highlights:**

- Cigarette smoking is strictly related to impairment of vascular repair mechanisms
- Reducing the exposure to toxicants in smoke, could reduce the harm to endothelium
- ENDS showed a reduced effect on endothelial cell migration compared to cigarette
- These data demonstrated the reduced toxicity of ENDS compared to cigarettes.

## Introduction

Cigarette smoking is a risk factor for many pathological conditions, including cardiovascular diseases (CVD) (Atlanta, 2014). According to the World Health Organization (WHO) more than 8 million people die each year due to the consumption of tobacco products, making smoking the leading cause of preventable deaths worldwide. Cigarette smoking is strictly related to endothelial dysfunction and structural damage to the endothelium (Lu et al., 2018), which leads to the impairment of vascular repair mechanisms, such as the inhibition of endothelial cell migration (Bernhard et al., 2003; Guarino et al., 2011; Newby et al., 1999). The ability to maintain the endothelium integrity is one of the most critical functions of the endothelial cells. Following endothelial injury there is an increased risk of developing vascular diseases such as atherosclerosis (Gotlieb and Lee, 1999), and an utmost need to rapidly restore the endothelial continuity. Forthwith, platelets and inflammatory cells adhere to the lesion to heal the wound triggering the migration and subsequent proliferation of medial smooth muscle cells in the neointima and thus, concurring to the development of occlusive vascular lesions (Packham and Mustard, 1986). Different studies investigated in vitro the detrimental effects of smoking-related harm to clarify the mechanisms and key events associated with the development of atherosclerosis, including cell migration inhibition (Fearon et al., 2013; Snajdar et al., 2001). Reducing the exposure to these toxicants therefore, may represent a possible strategy to reduce the harmful effect on the endothelium and, consequently, the effect of cigarette smoke on cardiovascular diseases. In particular, a study by Taylor et al. (Taylor et al., 2017) showed a significantly reduced inhibition of endothelial cell migration in vitro by electronic cigarette (e-cig) aerosol exposure when compared to cigarette (3R4F) smoke. In particular, E-cig is a non-combustible technology able to deliver nicotine to users with a lower toxicants content than smoke (Caruso M., 2021). Similarly, tobacco heated products (THPs) vaporize vegetable glycerin deposited on the tobacco with a working temperature within 350°C to provide users with an aerosol containing nicotine with an aroma similar to that of a cigarette, but with a lower content of combustion toxicants (Polosa et al., 2019). These products are generally referred to as electronic nicotine delivery systems (ENDS), and are often proposed as reduced-risk alternatives to the classic cigarette. However, the scientific debate is still open (Bals et al., 2019) and warrants further studies with particular regards to prolonged exposure to the aerosols released by ENDS.

The aim of the present study was to perform a multi-center replication study (ring study) to verify the results of Taylor and colleagues (Taylor et al., 2017) on the reduced ability of e-cigs aerosol to inhibit *in vitro* endothelial migration compared to cigarette smoke.

## Materials and methods

### Recruitment of Laboratories

International Laboratories experienced in maintaining HUVEC cultures were invited to participate in the inter-laboratory Replica study based on predefined criteria. An online questionnaire was administered to the international laboratories participating to the study that listed skills and knowledge pertaining to the core activities of this in vitro research to assess levels of proficiency in general and in relation to specific area of this research, including experience in assessments of endothelial cell migration and laboratory compliance with the Routine Analytical Cigarette-Smoking Machine — Definitions and Standard Conditions ISO3308:2012 (International Organization for Standardization 2018), European Good Laboratory Practice, and US Environmental Protection Agency Good Laboratory Practice Standards guidelines (Caruso et al., 2021).

The selected laboratories were provided with workshops, hands-on training, and on-site assessments of laboratory capacity and personnel expertise, with follow-up by virtual sessions, if necessary. Scientists received previous formal training in smoke and aerosol exposure procedures (Caruso et al., 2021) along with the standard operating procedures (SOPs) for use of smoking/vaping machines, cell-exposure systems and scratch wound assay. Four selected laboratories in academic establishments joined this study: one from each of Italy (LAB-A; leading center), Greece (LAB-B), USA (LAB-C), and Serbia (LAB-D).

### Harmonization Process

Laboratory protocols were harmonized across study sites with SOPs defined for each experimental step and use of the same cell lines, cell-exposure equipment, and methods to assess endpoints, as suggested by the Center for Open Science transparency and openness promotion guidelines (https://www.cos.io/initiatives/top-guidelines). A kick-off meeting held by LAB-A to introduce the SOPs and personnel training was provided as previously reported (Caruso et al., 2021). The SOPs for cell exposure to cigarette smoke and ENDS aerosol Aqueous Extract (AqE), cell culture and scratch wound assay were drawn up using the information contained in the original study (Taylor et al., 2017) and manufacturers’ instructions and adapted by the principal investigator sites according to laboratory capacity, equipment, and test products, ensuring they met the ISO3308:2012 guidelines (International Organization for Standardization 2018).

Detailed recording of technical data and deviation communication forms were collected by each laboratory partner by template spreadsheets (Microsoft Excel; version 16.43, 2011, Microsoft, Redmond, WA, USA) and shared with the leading laboratory, as previously stated (Caruso et al., 2021). To maximize assay standardization of cell growth and assessments, a list of key consumables was shared with all laboratories and these were obtained from the same lot when possible. A SOP was distributed for thawing, freezing, and subculturing of the cell line, including testing for mycoplasma contamination with the Plasmotest™ kit (InvivoGen, San Diego, CA, USA) before freezing the cells, to allow the laboratory partners to generate their own working cell bank.

### Original study

The study from Taylor and colleagues (Taylor et al., 2017) conducted a comparison study between the effects of two commercial e-cigarette products (Vype ePen and Vype eStick) and a scientific reference cigarette (3R4F) on endothelial migration in vitro. Here, we replicated the same study comparing the effects of three commercial ENDS (Vype ePen 3, Glo□ Pro and IQOS 3 DUO) and a scientific reference cigarette (1R6F). Vype eStick has been withdrawn from the market in many countries and the experimental 3R4F cigarette has been replaced by the 1R6F from the manufacturer (Center for Tobacco Reference Products, University of Kentucky). We measured the scratch wound area at different time points in order to quantify the migration over the time, and the percentage wound area of the initial wound and the time-by-time wound area, for each test product. Moreover, comparisons among each product response were reported.

### Test products

The following products were used for this study: (1) 1R6F reference cigarette (Center for Tobacco Reference Products, University of Kentucky); (2) Vype e-Pen 3 electronic cigarette (British American Tobacco); (3) Glo™ Pro (British American Tobacco); (4) IQOS Duo (Philip Morris International SA). The 1R6F cigarette has a tar yield of 29.1 mg/cigarette (Health Canada Intense [HCI] regime), and a nicotine content of 1.896 mg/cigarette. Before use, 1R6F cigarettes were conditioned for a minimum of 48 h at 22 ± 1 °C and 60 ± 3% relative humidity, according to ISO 3402:1999 (International Organization for Standardization, 1999). The Vype e-Pen 3 is an electronic cigarette with a “closed-modular” system consisting of two modules: a built-in 650 mAh rechargeable lithium battery section, and a replaceable “e-liquid” cartridge (“cartomizer”). The pods contain a reservoir of 2 ml of pre-filled liquid and the “Master Blend” (18 mg/ml nicotine) flavored variant was used for the experiment. Glo™ Pro and IQOS Duo are tobacco heating products (THPs). The THPs are devices that heat tobacco to generate a nicotine-containing aerosol with a tobacco taste inhaled by users. Glo□Pro device was used with “Ultramarine” Neostick, instead IQOS device was used with “Sienna Selection” Heatsticks. Each device was cleaned before and after use. All devices were fully charged and checked prior to being used.

### Preparation of aqueous aerosol extracts (AqE)

All the regimes used in this study were described in table 1. Whole smoke from 1R6F cigarette was generated on a LM1 smoking machine (Borgwaldt KC GmbH, Hamburg – Germany). 1R6F cigarette was smoked for 9 puffs, following the HCI puffing regime (55 ml, 2 s duration bell shape profile, puff every 30 s with filter vent blocked). Vype e-Pen 3 and the THPs were machine-puffed on a LM4E vaping machine (Borgwaldt KC GmbH, Hamburg – Germany). Vype ePen 3 was vaped for 10 puffs following the CRM81 regime (55 mL puff volume, drawn over 3 s, once every 30 s with square shape profile). IQOS Duo and GLO™ Pro were vaped using HCI regime without blocking the filter vents, for 8 (1 Neostick) and 12 (1 Heatstick) puffs respectively. The AqEs from 1R6F cigarette, e-cigarette and THPs were generated by bubbling through 20 ml of AqE capture media (VascuLife® media with added supplements and 0.1% of FBS). This procedure provided the AqE 100% stocks, which were diluted with appropriate volumes of VascuLife® media to produce the AqE concentrations for *in vitro* exposures. A range of concentration from 5 to 30% was used for the 1R6F cigarette. Instead, a range from 40 to 100% was used to test Vype ePen 3, IQOS Duo, and Glo □ Pro (Table 2).

**Table 1.** Puffing regime description for each product.

| Product | Puffing Regime | Puff Volume (ml) | Puff Frequency (sec) | Puff Duration (sec) | Puff Profile | Vent Blocking | Pre-activation (sec) |
| --- | --- | --- | --- | --- | --- | --- | --- |
| 1R6F | HCI | 55 | 30 | 2 | Bell | 100% | NA |
| Vype ePen3 | CRM81 | 55 | 30 | 3 | Square | NA | 0 |
| IQOS Duo | HCI | 55 | 30 | 2 | Bell | 0% | 30 |
| Glo□ Pro | HCI | 55 | 30 | 2 | Bell | 0% | 20 |

CRM81= Coresta Recommended Method n°81; HCI= Health Canada Intense; NA= Not Applicable

**Table 2.** AqE exposure concentration range for each test product.

| 1R6F Cigarette | Electronic Cigarette (ePen3) | THPs (IQOS Duo and Glo™ Pro) |
| --- | --- | --- |
| AqE (%) | AqE (%) | AqE (%) |
| 30 | 100 | 100 |
| 25 | 90 | 90 |
| 20 | 80 | 80 |
| 15 | 70 | 70 |
| 12.5 | 60 | 60 |
| 10 | 50 | 50 |
| 5 | 40 | 40 |

### Nicotine dosimetry

Nicotine dosimetry was carried out only by LAB-A on 100% diluted AqEs samples, collected after exposure for each product. A blank sample and six calibration standards, prepared in the same matrix at concentrations between 1-50 µg/ml (1, 2, 5, 10, 20, 50 µg/ml), were analyzed too. Aliquots of 0,1 ml of each sample and each calibration standards were transferred to vial with a 250 μl conical insert, added with nicotine-(methyl-d3) solution - used as internal standard at 10 µg/ml - and 0,1 ml of acetonitrile. Nicotine was determined by UPLC-ESI-TQD (Waters Acquity) with an Acquity UPLC® HSS T3 1.8 μm – 2.1×100mm column, operating in Multiple Reaction Monitoring (MRM) and positive ion mode. In Table 3 are reported ion transitions for nicotine and nicotine-(methyl-d3). Isocratic elution (80% water and 20% acetonitrile, both added at 0.1% with formic acid) was performed. The mass spectrometry settings were as follows: capillary energy at 2.5 kV, source temperature at 150°C, column temperature at 40 °C, desolvation temperature at 500 °C, desolvation gas at 1000 L/hr and cone gas at 100 L/hr. The injection volume was 1 µl.

**Table 3.** MRM mode: ion transitions (m/z) and relative cone and collision voltages.

| Analyte | MRM (m/z) | Cone (volts) | Collision energy (eV) |
| --- | --- | --- | --- |
| Nicotine | 163.0 → 117.0 | 40 | 25 |
|  | 163.0 → 132.0 | 40 | 15 |
| Nicotine-(methyl-d3) | 165.8 → 116.8 | 40 | 20 |
|  | 165.8 → 129.7 | 40 | 20 |

To calculate nicotine concentrations of each AqE dilution, a linear proportion was applied.

### Endothelial cell culture

Normal human umbilical vein endothelial cells (HUVECs; Lifeline Cell Technology, California, USA) were cultured as described by Taylor and colleagues (Taylor et al., 2017). Briefly, HUVECs have grown at 37°C and 5% CO2 in complete VascuLife VEGF Medium (Lifeline Cell Technology, California, USA), containing vascular endothelial growth factor (5 ng/mL), epidermal growth factor (5 ng/mL), basic fibroblast growth factor (5 ng/mL), insulin-like growth factor 1 (15 ng/mL), ascorbic acid (50 μg/mL), L-glutamine (10 mM), hydrocortisone hemisuccinate (1 μg/mL), heparin sulphate (0.75 units/mL), fetal bovine serum (FBS) (2%), penicillin (10,000 Units/mL), streptomycin (10,000 µg/mL), and amphotericin B (25 µg/mL). When the cells reached confluence, they were detached with trypsin-EDTA solution (0.05%) and replated in new flasks or into 24-well plates and used in experiments. Cultured HUVECs maintain their normal appearance for 15 population doublings. Then, we discarded HUVECs after 4 passage cycles.

### Endothelial cell scratch wound assay

HUVECs were seeded in 24-well plates at a density of 2 × 10^5^ cells/well in complete VascuLife VEGF Medium, and incubated at 37 °C and 5% CO2 until they reached the total confluency (24–48 h prior to performing the assay). When the HUVECs were ready to perform the scratch wound assay, the complete VascuLife VEGF Medium was replaced with AqE capture media (VascuLife® media containing 0.1% FBS), then cells were incubated for 6 h. Linear scratch wounds were created manually by using sterile 10 μL pipette tips. Immediately after wounding, the medium with detached cells was removed, and a washing step with PBS was performed. Next, cells were exposed with 1 ml of each test product AqE in triplicate. A negative control with not exposed AqE capture media and a positive control with cytochalasin D (2 μM) were entered for each plate. One laboratory (LAB-A) used the Operetta CLS™ high-content analysis system to read the experimental plates by using a ×5 objective to acquire the images. Instead, the other laboratories (LAB-B, LAB-C, and LAB-D) incubated the plates into an incubator at 37 °C and 5% CO2. Then the cells were taken out of the incubator at established time-points ranging from 0 to 48 hours (T0 - T48) and placed under a microscope with ×5 objective. Two pictures per well of fixed positions in the wounds were taken with a digital camera mounted on the microscope.

### Statistical analysis

The scratch wound area (μm^2^) was measured at each time point using the open-source software imageJ/Fiji®. In order to quantify the migration over the time, the percentage wound area of the initial wound area was calculated by the following formula: A(Tn)%= A(Tn)/A(T0)*100, where A(Tn) is the wound area at time Tn and A(T0) is its initial area. Comparisons among the tested concentrations were analyzed by fitting a repeated measure mixed model followed by Dunnet’s test to perform multiple comparisons with the untreated cellular response. Moreover, comparisons among each product response were analyzed by fitting a repeated measure mixed model followed by Tuckey’s test adjustment for multiple comparisons. Data were expressed as mean ± standard error (SE). All analyses were considered significant with a p-value of less than 5 %. We analyzed and plotted the results using GraphPad Prism version 8 (GraphPad Software, San Diego, California, USA, www.graphpad.com).

## Results

### AqE nicotine dosimetry

Analysis on 100% diluted AqE samples showed nicotine concentrations of 12.8 μg/mL for 1R6F, 4.2 μg/mL for Vype ePen, 8.4 μg/mL for IQOS and 4.5 μg/mL for Glo, respectively. The calculated concentrations of nicotine for each tested AqE dilution were reported in table 4.

**Table 4.**
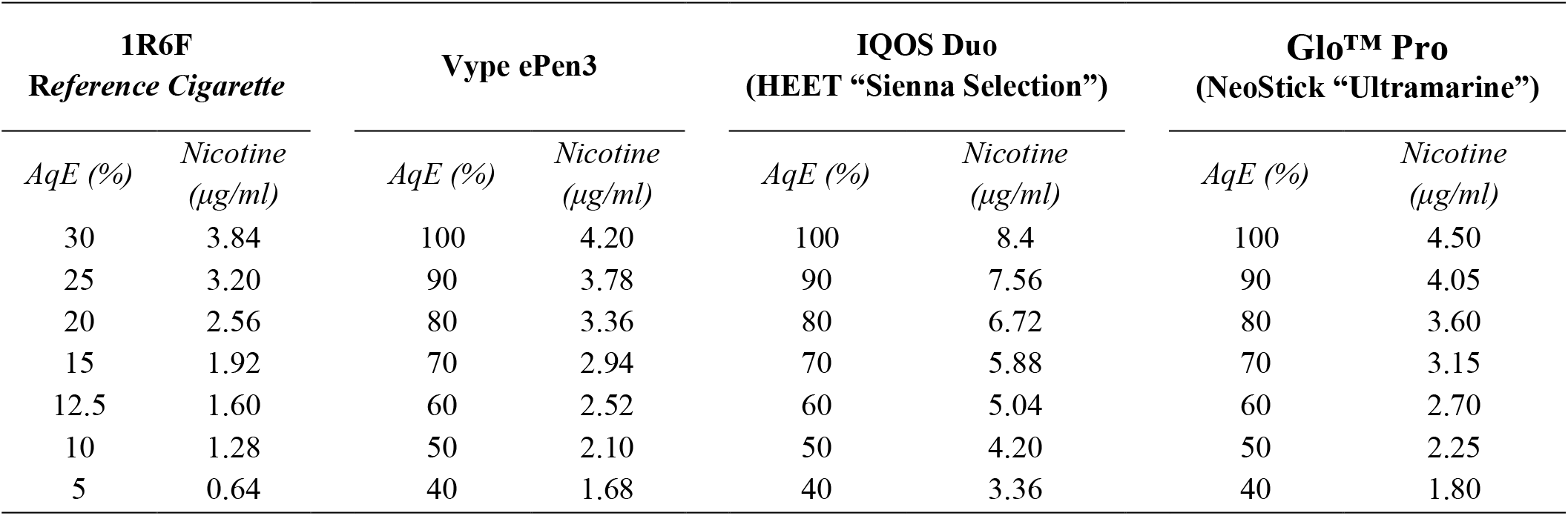
Nicotine concentration calculated for each AqE dilution.

### HUVEC cell migration baseline

For each scratch wound assay a negative control with AqE capture media, and a positive control with cytochalasin D (2 μM) were evaluated to determine the baseline and the maximal inhibition of HUVEC migration, respectively. The untreated HUVECs incubated with AqE capture media showed a time-dependent closure of wound area (Fig. 1). We observed a mean wound area percentage under 20% starting from T-20 (A(T20)%= 16.33 ± 1.85 %) until a complete wound closure at T-48 (A(T48)%= 0.91 ± 0.5 %). However, the incubation with cytochalasin D totally inhibited the closure of the wound during the 48 hours exposure period (Fig. 1). Representative images of wound healing after AqE capture medium and cytochalasin D treatments were shown in figure 2.

**Figure 1.**
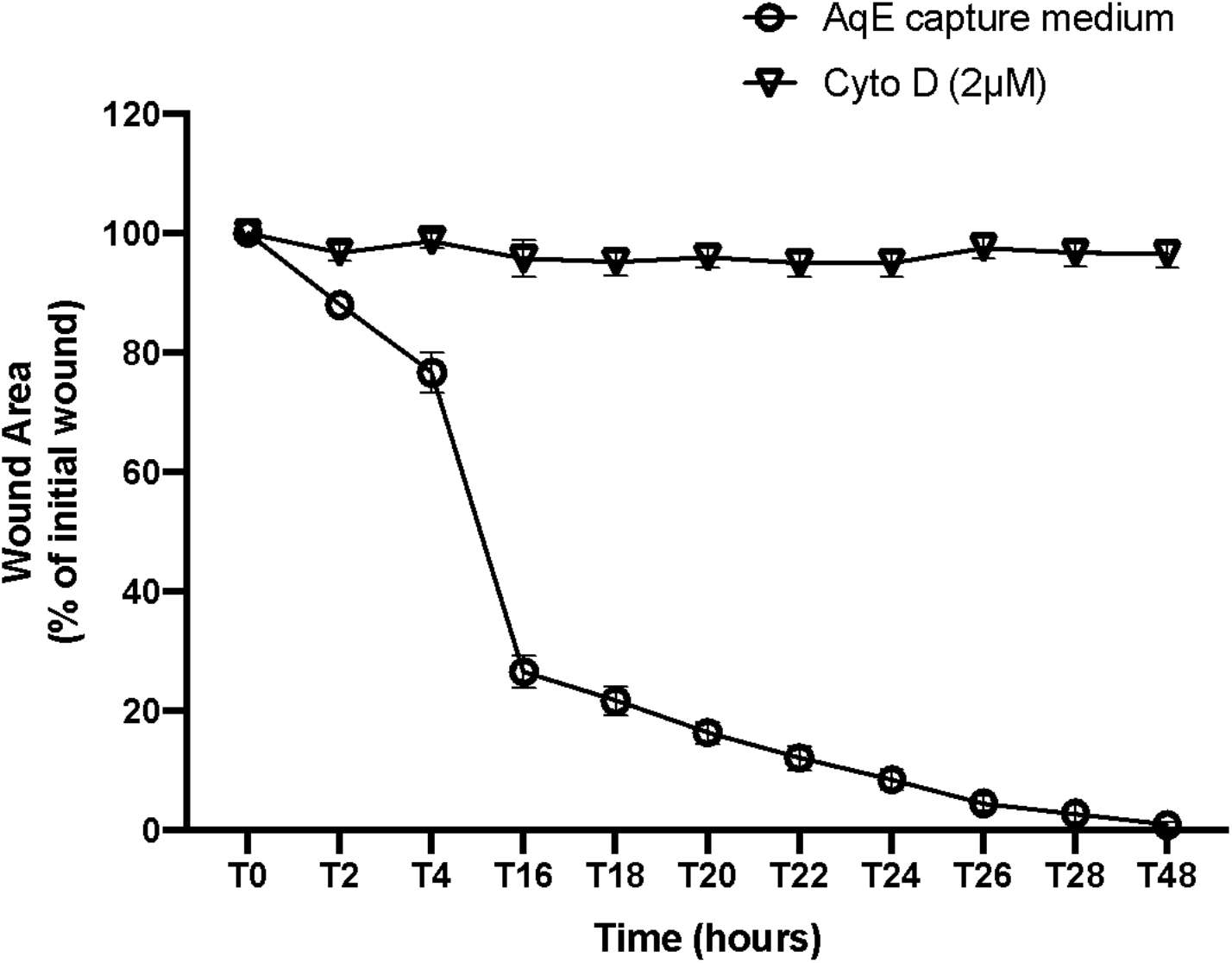
The incubation of HUVECs with AqE capture media allowed the closure of wound with a time-dependent reduction of wound area over 48 h. Cytochalasin D (2 μM) inhibited HUVECs migration. Data were showed as mean ± standard error (SE) of wound area (μm^2^) percentage from duplicate wells of 4 independent experiments.

**Figure 2.**
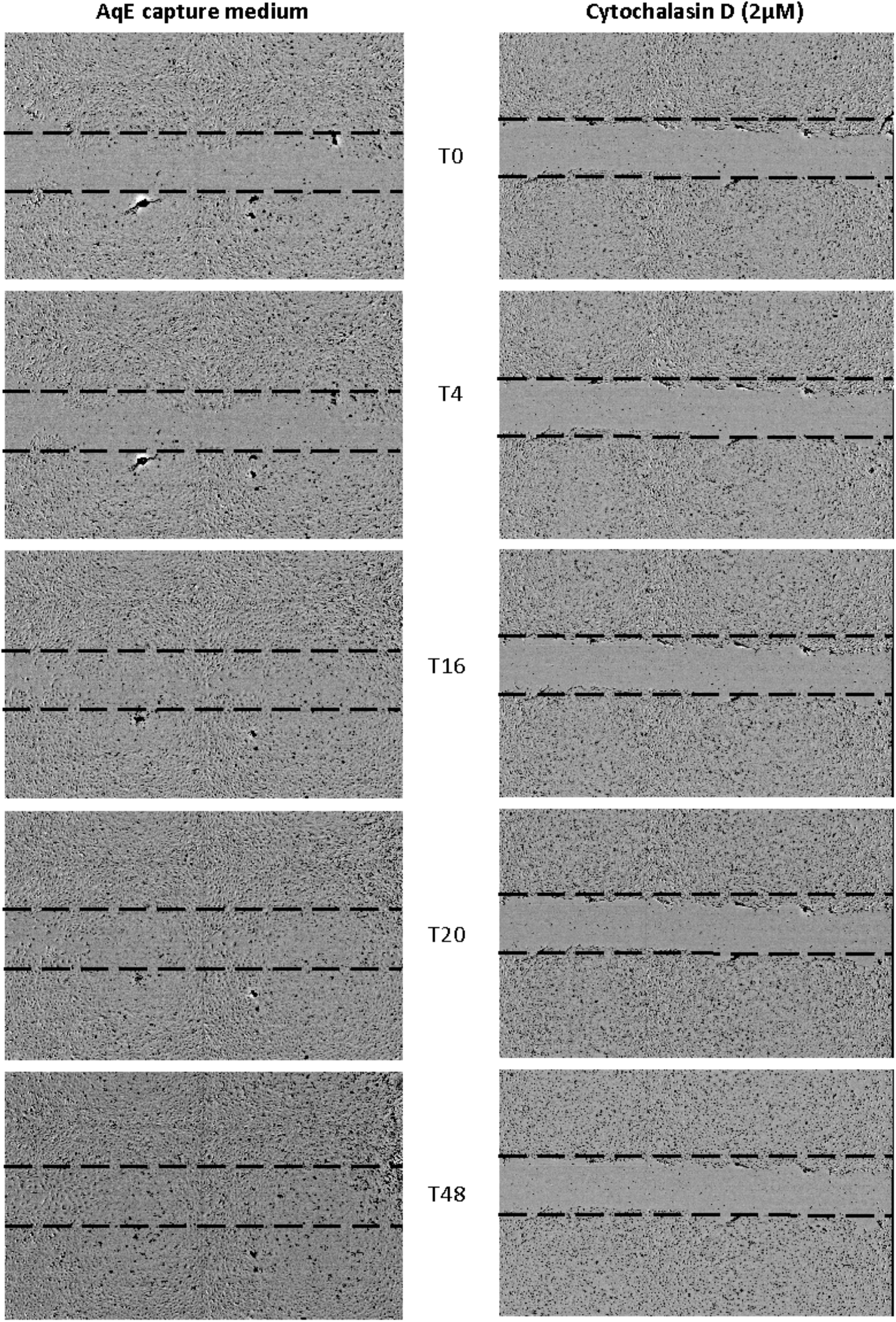
Representative images of wound healing in the presence of AqE capture medium and cytochalasin D at T0, T4, T16, T20, and T48.

### HUVEC migration after exposure to cigarette smoke AqE

Cigarette smoke AqE on HUVEC migration was assessed across concentrations ranging from 5 to 30% 1R6F AqE, which represent exposure to 0.64–3.84 μg/mL of nicotine. Exposure to 1R6F AqEs inhibited HUVEC migration in a concentration-dependent manner, confirming findings by Taylor and colleagues (Taylor et al., 2017). In particular, we observed a significant inhibition of wound area closure for concentrations ranging from 12.5% to 30% compared to AqE capture media treatment (p<0.05). Only the 5% 1R6F AqE allows a complete wound closure at 48h. Complete inhibition of endothelial migration was observed when the cells were exposed to 25% and 30% 1R6F AqE (Fig. 3).

**Figure 3.**
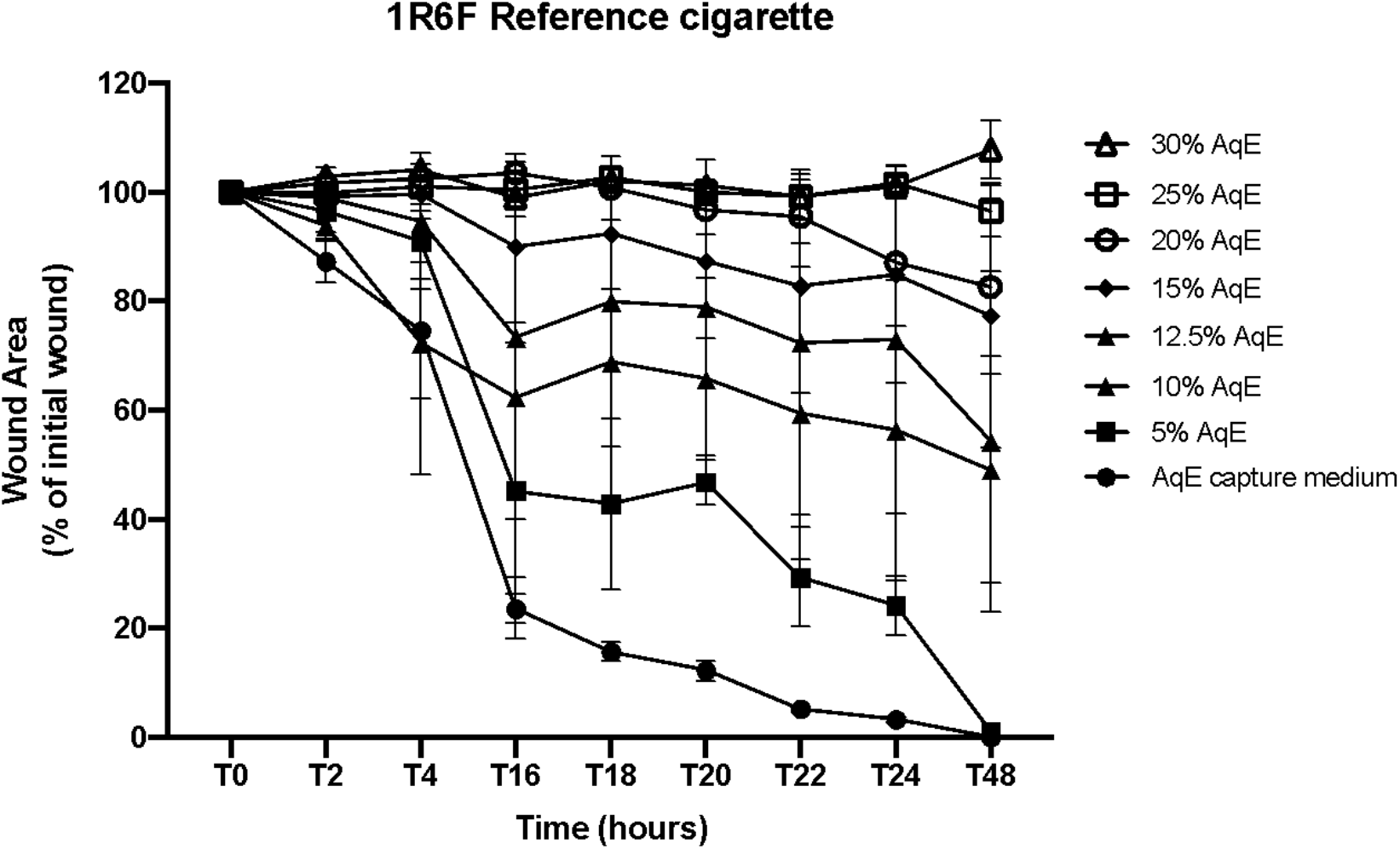
The HUVEC wound healing was inhibited by 1R6F AqE in a concentration-dependent manner. Data are reported as mean ± standard error (SE) of wound area (μm^2^) percentage from triplicate wells 4 independent experiments.

**Figure 4.**
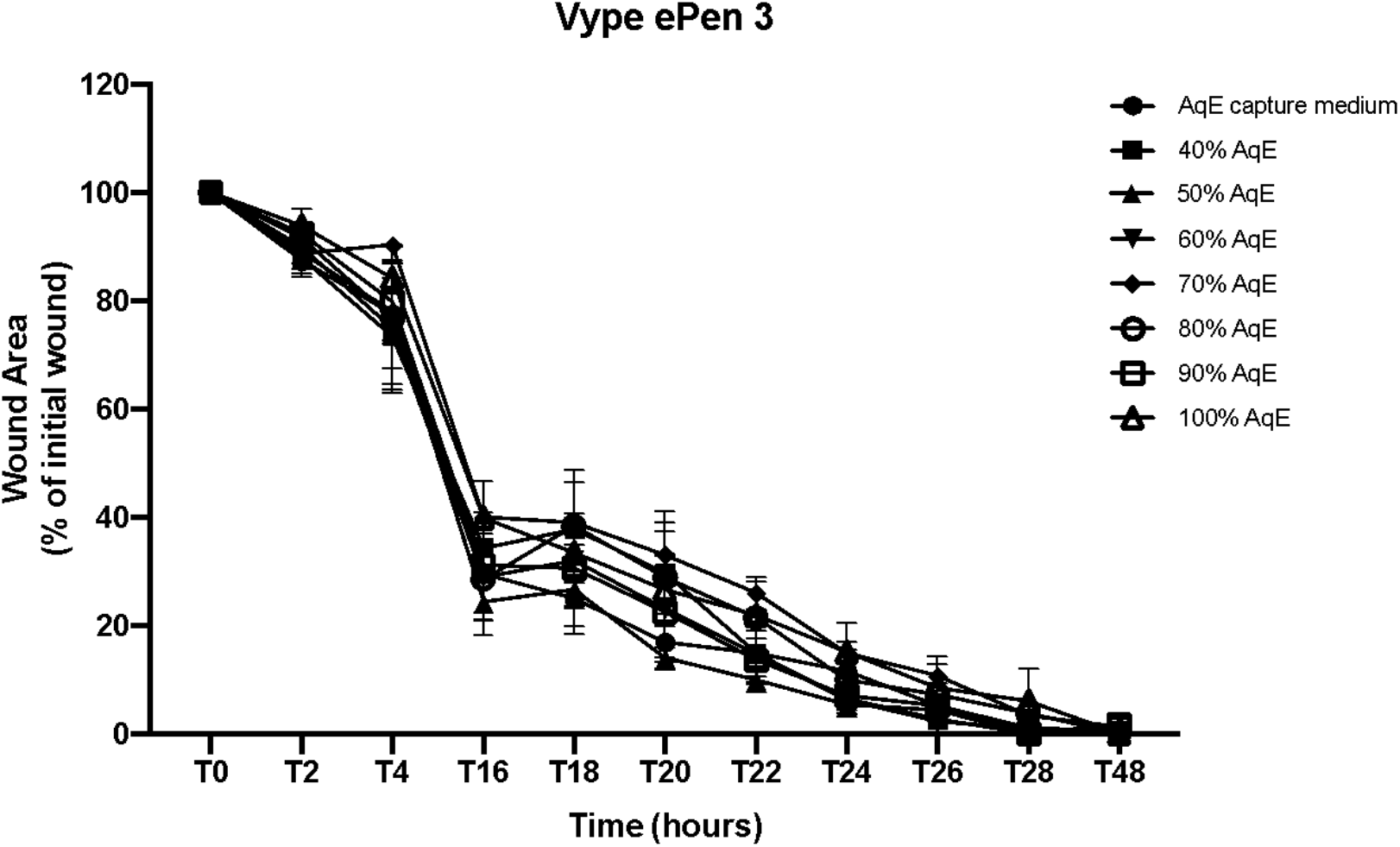
Vype ePen3 AqE did not inhibit HUVEC migration. Data are reported as mean wound widths as mean ± standard error (SE) of wound area (μm^2^) percentage from triplicate wells of 4 independent experiments.

### HUVEC migration after exposure to e-cigarette AqE

We tested Vype ePen 3 AqE with a range from 40 to 100%. These AqE concentrations contained a 1.68–4.2 μg/mL nicotine (Table 4). All the Vype ePen 3 AqE concentrations did not produce any significant reduction of wound area closure compared to AqE capture media treatment (Fig.4).

### HUVEC migration after exposure to THP AqEs

Both IQOS Duo and Glo™ Pro AqEs were tested with concentration ranging from 40 to 100%. This AqE concentration range contained 3.36–8.4 μg/mL nicotine for the IQOS Duo and 1.8–4.5 μg/mL nicotine for the Glo™ Pro. IQOS Duo AqEs with concentrations ranging from 40% to 80% did not affect the HUVEC migration compared to AqE capture media (p> 0.05). Instead, significant differences were observed for 90% (p= 0.009) and 100 % (p= 0.002) IQOS Duo AqEs when compared to AqE capture media (Fig. 5), despite the complete closure of wound at T48. All the Glo™ Pro AqE concentrations did not reduce the HUVEC migration compared to AqE capture media (Fig. 6).

**Figure 5.**
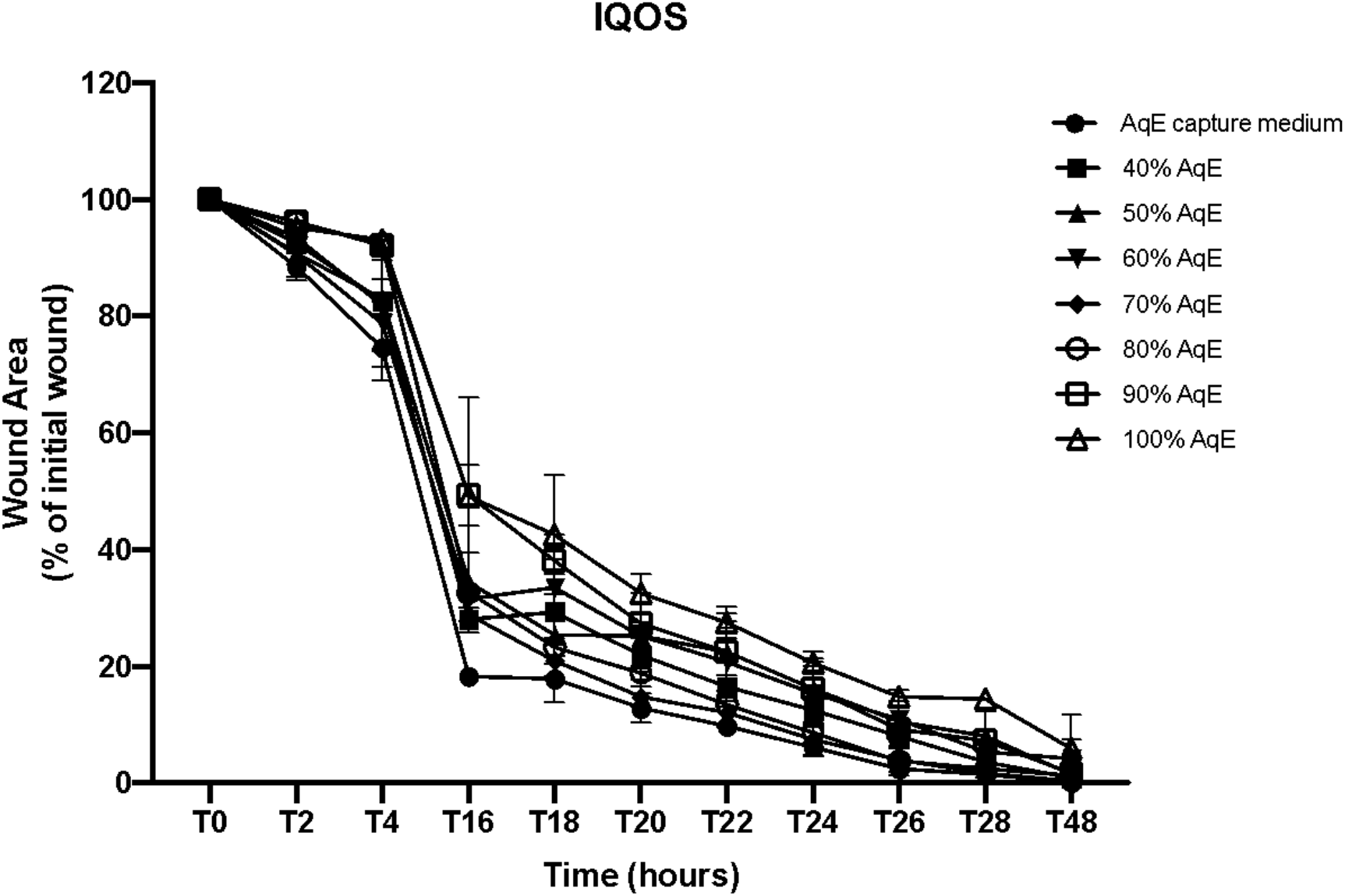
IQOS Duo AqE effect on HUVEC migration. Data are reported as mean wound widths as mean ± standard error (SE) of wound area (μm^2^) percentage from triplicate wells of 4 independent experiments.

**Figure 6.**
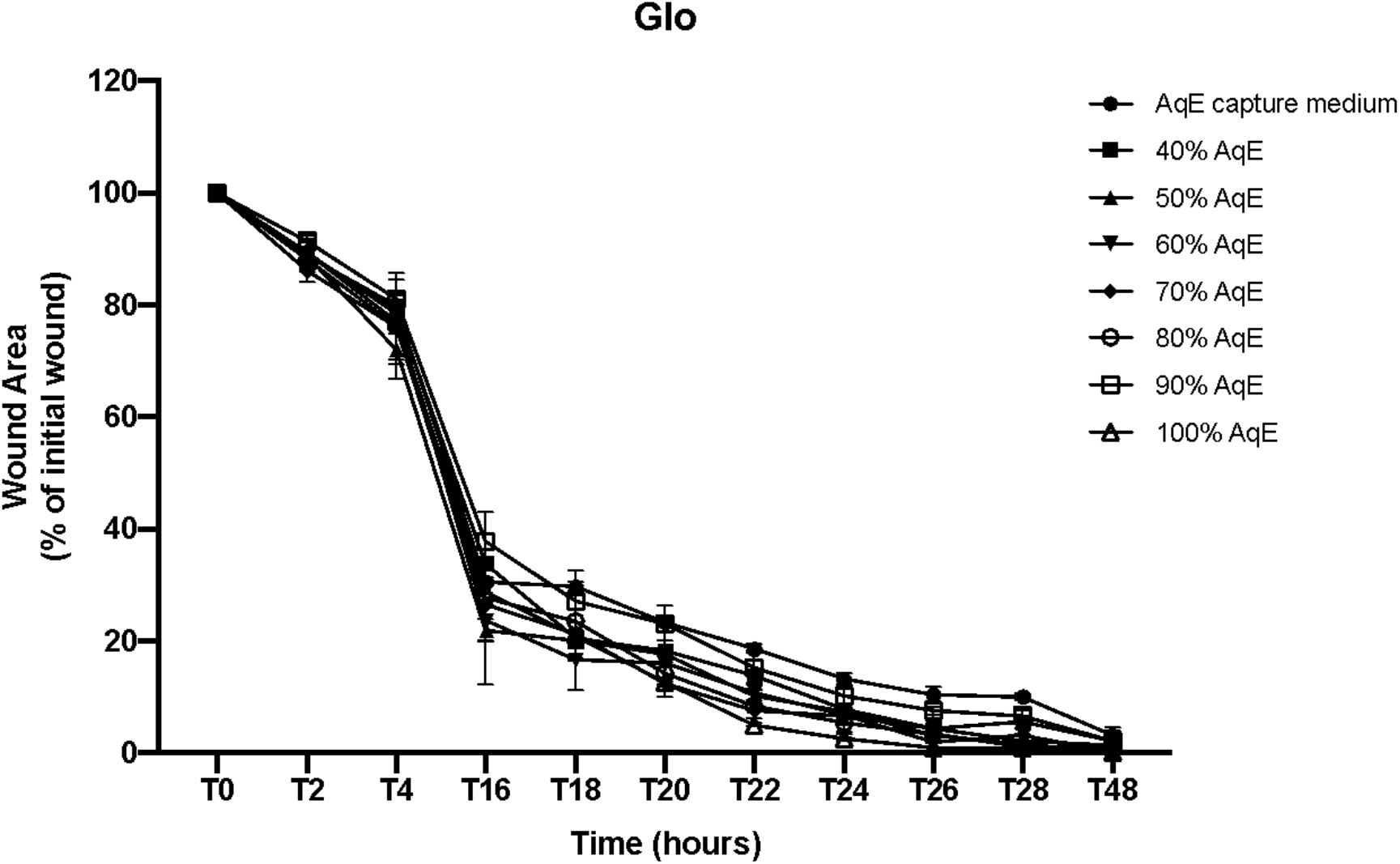
Glo □ Pro AqE effect on HUVEC migration. Data are reported as mean wound widths as mean ± standard error (SE) of wound area (μm^2^) percentage from triplicate wells of 4 independent experiments.

### Comparison among Cigarette, ePen, IQOS and Glo exposures

A comparison of HUVEC migration was made among all the tested products at the higher concentration (Fig. 7). Significant differences were observed for Vype ePen 3 100%, IQOS Duo 100%, and Glo™ Pro 100% AqEs compared to 1R6F 30% AqE (p< 0.0001). Moreover, a slight significant difference was shown between IQOS Duo 100% and Glo™ Pro 100% AqEs (p= 0.039). No differences were observed between Vype ePen 3 100% and THPs 100% AqEs (p> 0.05). Even comparing the migration of endothelial cells between all alternative products with equal or greater amounts of nicotine released (Fig. 8) by 1R6F cigarette at higher concentration (3.84 μg/ml), we observed significant differences. Finally, comparison of the migration of endothelial cells exposed to the maximal concentration of each product AqE (Fig. 6) highlighted a significant difference between ENDS and 1R6F cigarette AqE.

**Figure 7.**
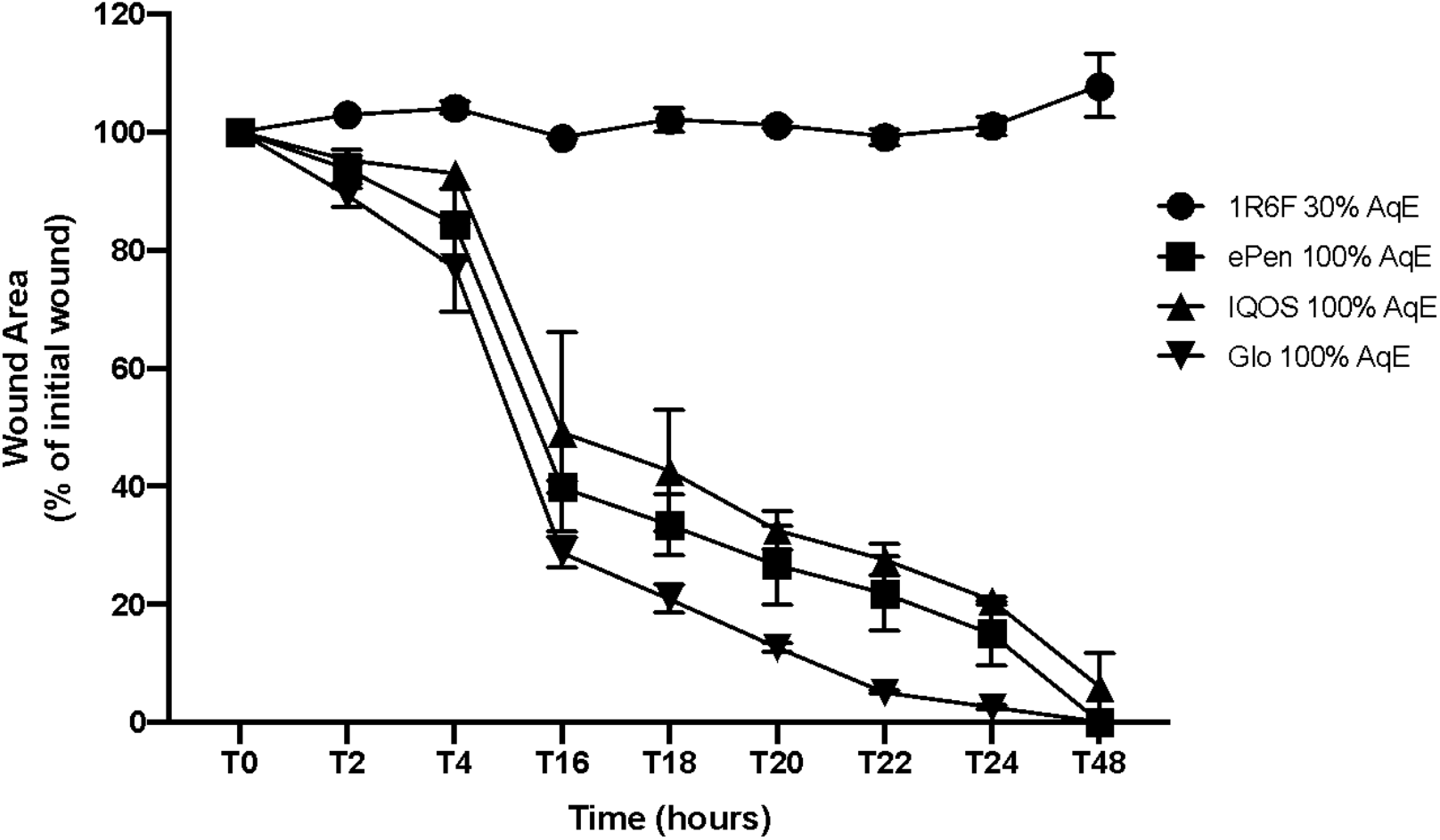
Comparisons among all the test products at the higher concentrations. Data are reported as mean wound widths as mean ± standard error (SE) of wound area (μm^2^) percentage from triplicate wells of 4 independent experiments.

**Figure 8.**
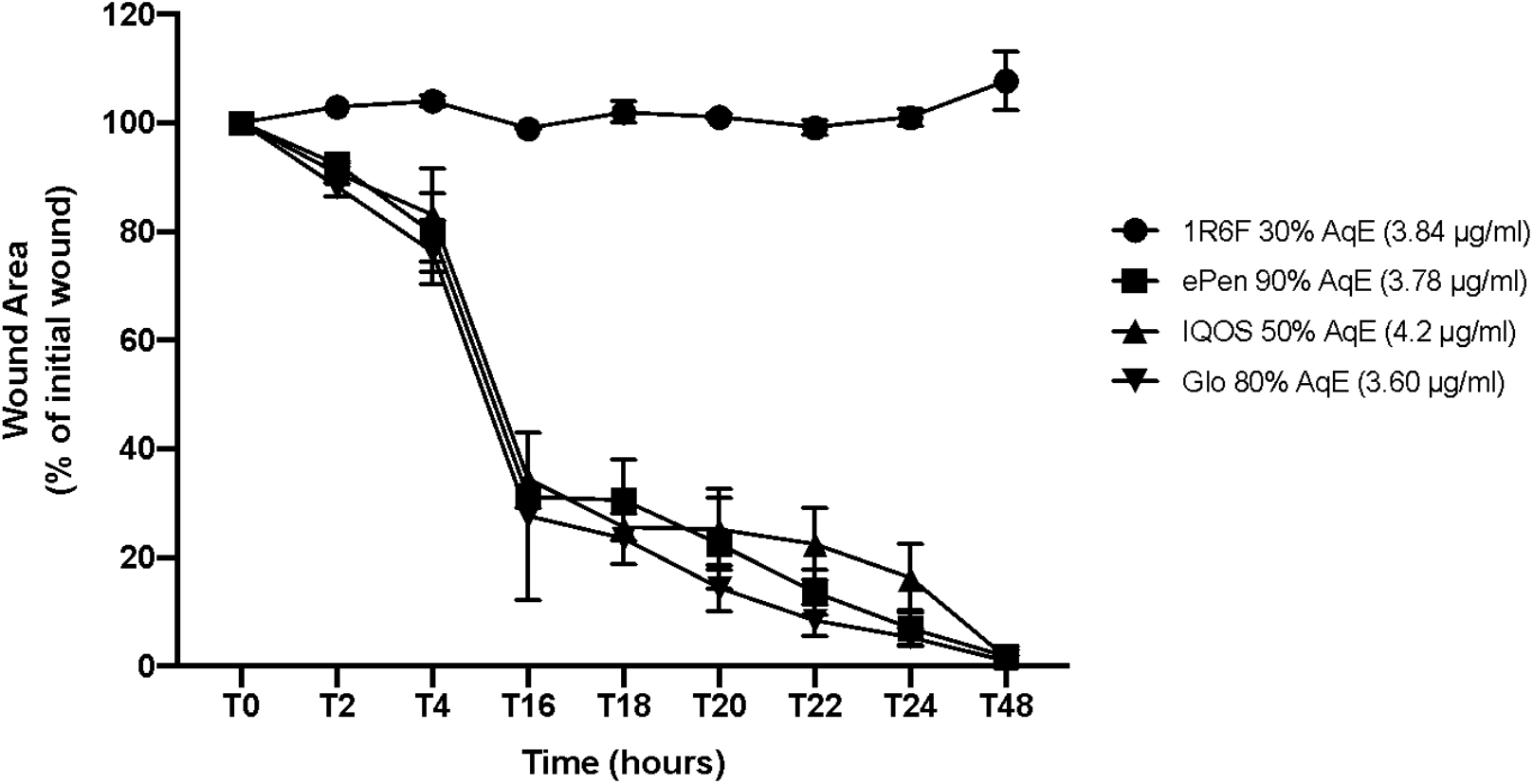
Comparisons among all the test products at similar nicotine concentrations. Data are reported as mean wound widths as mean ± standard error (SE) of wound area (μm^2^) percentage from triplicate wells of 4 independent experiments.

## Discussion

The detrimental effects of tobacco smoking to the cardiovascular system are well established (Atlanta, 2014), although the exact molecular mechanisms are yet to be fully defined. Taylor and colleagues (Taylor et al., 2017) investigated the ability of both cigarette smoke and e-cigarette aerosol to affect vascular endothelial cells wound healing, one of the key processes in atherosclerotic disease initiation and progression in the cardiovascular system. The authors used an *in vitro* model of endothelial cells exposed to AqE, the water-soluble fraction of smoke or e-cigarette aerosols, to resemble the in vivo exposure of endothelial cells to toxicants. The results from Taylor and colleagues demonstrated that cigarette smoke AqE resulted in a concentration-dependent inhibitory effect on endothelial cell migration, but no significant inhibition of endothelial cell migration following exposure to e-cigarette AqEs was observed. Furthermore, they hypothesized that chemical species present in the smoke AqE were responsible for the inhibition of endothelial cell migration and that these chemicals are absent, or present in insufficient concentrations, in the e-cigarette AqEs to elicit any significant response in the wound healing assay. In the present study, we replicated the paper by Taylor and colleagues in a multicenter study. The choice of a multi-center approach was used to verify the robustness and reliability of the results obtained in the original study. However, some methodological issues need to be clarified to fully explain our results. In particular, we used an updated version of the electronic cigarette device compared to that used in the original study, the Vype ePen 3 in place of the Vype ePen device, and we included two heated tobacco devices, IQOS Duo and Glo™ Pro. Additionally, the tobacco cigarette used in our study (1R6F) was different from that used in the original study (3R4F) since this has been out of production in recent years and the manufacturer recommends the 1R6F as a replacement for the 3R4F. The results on the ability of endothelial cells to heal the wound following exposure to cigarette smoke and e-cigarette aerosol from the four laboratories involved in this ring-study confirmed the results reported in the original study, despite the difference in the method used to assess the size of the wound at each time-point. In particular, Taylor and colleagues used the wound width as the measurement parameter for wound healing at each time-point and for each treatment. The wound width measurements were automatically acquired by using the IncuCyte live-cell imaging system. In our multicenter study, one laboratory used an automated live-cell imaging system to acquire the scratch images whereas the other three participating laboratories obtained scratch imaging manually with a digital camera mounted on the microscope. All the centers used the same software (imageJ/Fiji®) to measure the wound area. Furthermore, we found a higher nicotine concentration in 1R6F AqE compared to that reported by Taylor and colleagues. This difference could be due to the different smoking machines used in our study. Despite the difference in nicotine concentrations in cigarette AqEs, the exposure concentration range for the e-cigarette AqE included an equivalent nicotine dose to the 30% 1R6F AqE. As reported in a previous paper, we observed no significant variations in endothelial migration rates following exposure to different concentrations of Vype ePen 3 AqE, differently from what observed with the1R6F smoke AqEs which showed a consistent concentration-dependent effect. The different migration response of endothelial cells is significant only for the cigarette AqE thus supporting the results by Taylor and colleagues, namely that the inhibitory effect on cell migration and wound repair must be exerted by other chemicals contained in the smoke and absent, or scarcely present, in the ENDS aerosol, and not by nicotine. On the other hand, different studies have demonstrated that nicotine may induce migration and proliferation of vascular cells by binding to specific nicotinic acetylcholine receptors (Liu and Yeh, 2014; Park et al., 2008) and by increasing proangiogenic VEGF (Kim et al., 2017). Another conceivable hypothesis on the detrimental effects of smoke on migration and proliferation of endothelial cells could be the high quantity of oxidative species present in cigarette smoke. These chemicals induce endothelial dysfunction through oxidative damage to membrane constituents, mitochondria and DNA (Kannan and Jain, 2000) inducing both apoptosis and necrosis (Messner et al., 2012), as a results from oxidative damage to mitochondria or DNA. Differently from smoke, aerosol from e-cigs contains a substantially reduced number and quantity of such chemical species (Goniewicz et al., 2014; Taylor et al., 2016) suggesting a possible explanation for e-cig reduced endothelial cell toxicity.

We also studied vascular wound repair following the aerosol exposure of two commercially available THPs, IQOS duo and Glo™ Pro. These set of experiments showed no significant variation in wound healing for all the tested concentrations of Glo AqE. However, we only observed a slight inhibitory effect in wound closure for the 90% and 100% IQOS AqEs compared to media control, however significantly less than that observed with the 30% dilution of 1R6F AqE. Surprisingly, there have been few *in vitro* studies on THPs and their effect on vascular endothelial wound repair. To this regard, in 2017 Breheny and colleagues performed an *in vitro* screening of a commercial and prototype THPs, showing an inhibitory effect on HUVEC wound healing for the prototype THP at higher concentrations. Interestingly, this effect was not as evident as the inhibition observed with the AqE from cigarette (Breheny et al., 2017). In another study by British American Tobacco, the AqE from THP1.0 (GLO™ heating device with KENT Neosticks) did not affect HUVEC wound closure. Consistently, the highest concentration of 100 %, the relative density of wound closure has a similar trend to media control (Bishop et al., 2020). Finally, in 2015 van der Toorn and colleagues showed that AqE from Tobacco Heating System 2.2 (THS2.2, commercialized under the brand name IQOS®) affect the integrity of human coronary artery endothelial cell (HCAEC) monolayer but in a reduced manner compared to AqE from cigarettes (van der Toorn et al., 2015).

In conclusion, our data confirmed the results from Taylor and colleagues (Taylor et al., 2017) showing that AqEs of a commercially available e-cigarette (Vype ePen 3) does not induce the inhibition of endothelial cell migration *in vitro* as compared to cigarette smoke AqE. We additionally demonstrated a product-specific response on HUVEC migration for THPs. Particularly, it appears that IQOS possesses the potential to induce adverse cellular effects on the cardiovascular system, however such effects are much less pronounced than those observed with cigarette smoke. Our results provide useful scientific informations in support of the decision-making process of regulating these products in order to develop evidence-based harm reduction strategies and policy decisions by governments.

## Abbreviations

AqE: Aqueous Extract
ENDS: electronic nicotine delivery systems
THPs: tobacco heating products
E-cigs: Electronic Cigarettes
HCS: High Content Screening
ISO: International Organization for Standardization
HCI: Health Canada Intensive
CRM81: CORESTA Recommended Method n. 81

## Author contributions

Conceptualization: M.C., G.L.V., R.Po.; Methodology: M.C., G.L.V.; Formal analysis and investigation: R.E., A.D., K.M., S.B., A.A., G.K., R.Pu., A.I., A.C., P.Z.; Writing—original draft preparation: M.C., G.L.V., R.E.; Writing—review and editing: M.C., S.R., K.P., A.G., V.V., M.F., R.Po., G.L.V.; Supervision: M.C., G.L.V.

